# Protein Complex Structure Prediction Powered by Multiple Sequence Alignments of Interologs from Multiple Taxonomic Ranks and AlphaFold2

**DOI:** 10.1101/2021.12.21.473437

**Authors:** Yunda Si, Chengfei Yan

## Abstract

AlphaFold2 is expected to be able to predict protein complex structures as long as a multiple sequence alignment (MSA) of the interologs of the target protein-protein interaction (PPI) can be provided. In this study, a simplified phylogeny-based approach was applied to generate the MSA of interologs, which was then used as the input to AlphaFold2 for protein complex structure prediction. Extensively benchmarked this protocol on non-redundant PPI dataset including 107 bacterial PPIs and 442 eukaryotic PPIs, we show complex structures of 79.5% of the bacterial PPIs and 49.8% of the eukaryotic PPIs can be successfully predicted, which yielded significantly better performance than the application of MSA of interologs prepared by two existing approaches. Considering PPIs may not be conserved in species with long evolutionary distances, we further restricted interologs in the MSA to different taxonomic ranks of the species of the target PPI in protein complex structure prediction. We found the success rates can be increased to 87.9% for the bacterial PPIs and 56.3% for the eukaryotic PPIs if interologs in the MSA are restricted to a specific taxonomic rank of the species of each target PPI. Finally, we show the optimal taxonomic ranks for protein complex structure prediction can be selected with the application of the predicted TM-scores of the output models.

## Introduction

Proteins often facilitate their functions through interacting with other proteins. Therefore, the knowledge of protein complex structures formed by interacting proteins is valuable for mechanistic investigation of life processes and therapeutic development[1–4]. However, by far, only a small fraction of protein-protein interactions (PPIs) have their experimental complex structures available in the Protein Data Bank (PDB)[5], which is mainly because determining the complex structures through experimental techniques like X-ray crystallography, cryogenic electron microscopy (cryo-EM), etc. is time consuming and expensive[6]. Comparing with experimental structural determination, computational prediction is much more efficient and cheaper. Therefore, computational prediction of protein complex structures has become an important scientific problem for decades. The existing computational methods to predict protein complex structures can be grouped into two categories: protein-protein docking approach and template-based approach[7]. Protein-protein docking approach first applies sampling algorithms to generate a large number of possible binding modes between interacting proteins, then the produced binding modes are ranked using scoring functions to identify the most probable binding mode[8, 9]. Protein-protein docking approach has two major limitations. First, protein-protein docking generally uses the structure of each monomeric protein as the input to generate the possible binding modes, however, the monomeric structures of the interacting proteins are not always available. Second, proteins when forming complex structures often have conformational changes, which makes the configurational space of PPIs extremely large, therefore, the sampling algorithms in protein-protein docking is not always able to successfully generate the correct binding modes, or even near-native binding modes can be generated in the sampling stage, the scoring functions have difficulties to successfully identify these modes[10]. The template-based approach is based on the fact that protein-protein interologs often have similar complex structures, which applies the experimental complex structures of interologs of the target PPIs as the templates to build its complex structure[11]. The template-based approach can be very accurate when high quality templates are available. However, at the current stage, we still lack good templates for most of PPIs.

Recently, AlphaFold2 (AF2) has achieved unprecedented breakthrough in protein monomeric structure prediction[12, 13]. After that, in the work of RoseTTAFold[14], a model based on a similar deep learning architecture, the authors further pointed out that this deep learning architecture can also be used to predict the complex structure of intra-species PPIs. As long as a multiple sequence alignment (MSA) of a certain number of interologs of the target PPIs is provided, the deep learning architecture is able to infer the intra- and inter-protein evolutionary and coevolutionary information to build the protein complex structure. However, it is not trivial to generate the MSA of interologs, since a protein often have multiple homologous proteins including orthologs and paralogs in each species, and the interaction relationships between the two groups of homologous proteins of the interacting proteins are often unknown[15, 16]. In the work of RoseTTAFold, the interaction relationships are inferred based on their genomic distances, in which the sequences with similar Uniprot accession codes (i.e. shorter genomic distances) are paired[14]. This protocol is based on the theory of operon, which states that in prokaryotes, proteins with related functions are usually encoded together within the genome[17, 18]. However, operons are mainly observed in prokaryotes, therefore, this protocol may not work well for PPIs of eukaryotes. Besides, even prokaryotes also have a large number of PPIs in which the interacting proteins are not co-localized on the genomes, for which this protocol may also do not work well. Apart from the genomic distance-based approach, in the work of ComplexContact[19], a web server for predicting inter-protein residue-residue contacts, the authors applied a phylogeny-based approach to generate the MSA of interologs, in which the homologous proteins of each target interacting protein in each specific species were first descendingly sorted according to their sequence similarities to the target protein, and then the proteins with the same rankings within each species were paired to form MSA of interologs. However, whether the interaction relationships between proteins in the two homologous protein groups can be accurately inferred by this phylogeny-based approach has not been extensively validated. Besides, Gueudré *et al.* and Bitbol *et al.* also developed computational approaches respectively to infer protein-protein interologs through maximizing the coevolution signal between interacting homologous protein families[20–22]. However, the expensive computational cost of these approaches hinders their large-scale applications.

In this study, we employed a simplified version of the phylogeny-based approach to generate the MSA of interologs, in which only the sequences with the highest sequence similarity to the interacting proteins among the homologous proteins in each species were paired to form the (putative) interologs, for these proteins are more likely to be orthologs of the target interacting proteins, thus they have higher probability to interact with each other and are more likely to share a similar binding mode with the target interacting proteins. Different from the phylogeny-based approach of ComplexContact, we did not include the protein pairs with lower rankings in the MSA generation for two reasons: First, the lower ranking proteins are more likely to be paralogs of the interacting proteins, and the interaction of paralogs is often not as conserved as orthologs; second, even though the interaction of parologs is conserved, the accuracy of the inferred interaction relationships between the paralogs cannot be guaranteed. The MSA of interologs prepared by this simplified phylogeny approach was then employed as the input to the deep learning model of AF2 to predict the complex structure of interacting proteins, in which no template information was used to assist the protein complex structure prediction. This protocol was applied to predict complex structures for non-redundant intra-species PPIs including 107 bacterial PPIs and 442 eukaryotic PPIs. The result shows that we can successfully predict complex structures (DockQ≥0.23) for 79.5% of the bacterial PPIs and 49.8% of the eukaryotic PPIs with the application of this protocol. For the purpose of comparison, we also employed the MSAs of interologs prepared by the genomic distance-based approach of RoseTTAFold and by the phylogeny-based approach of ComplexContact as the inputs to AF2 to predict protein complex structures for the same dataset respectively. The result shows with the application of the genomic distance-based MSA in the protein complex structure prediction, we can successfully predict complex structures for only 56.1% of the bacterial PPIs and 17.6% of the eukaryotic PPIs; and with the application of the phylogeny-based MSA of ComplexContact, we can successfully predict complex structures for 78.5% of the bacterial PPIs and 47.0% of the eukaryotic PPIs, from which we can conclude that both the simplified phylogeny-based approach and the phylogeny-based approach of ComplexContact are significantly more effective than the genomic distance-based approach in obtaining the MSA of interologs for protein complex structure prediction, but the inclusion of the lower ranking protein pairs in the phylogeny-based MSA of interologs can even attenuate the protein complex structure prediction performance. Since PPIs are generally not as conserved as monomeric proteins in the process of evolution, a PPI may not be maintained in another species with very long evolutionary distance, or even though the PPI is maintained, the complex structure of the interolog can be different[23, 24]. Including non-conserved interologs in the MSA may attenuate the prediction performance as well. To verify this assumption, we further applied MSAs in which the interologs are from species within different taxonomic ranks (domain, kingdom, phylum, class and order) of the species of the target PPI in the complex structure prediction. We found that for both the bacterial and the eukaryotic PPI datasets, higher prediction accuracies for a significant number of PPIs can be achieved when restricting interologs in the MSA to each taxonomic rank. If the optimal taxonomic rank for each target can be chosen in the MSA generation, the success rates for the protein complex structure prediction can be increased to 87.9% for the bacterial PPIs and 56.3% for the eukaryotic PPIs. Finally, we show that the predicted TM-scores (pTM) of the generated models can be used to select the optimal taxonomic ranks for protein complex structure prediction.

## Materials and Methods

The protein complex structure prediction and evaluation protocols are overviewed in this section. For a more detailed description of the preparation of the test dataset and the implementation of the prediction and evaluation protocols, please refer the Supplemental Methods section provided in the Supplemental PDF.

### 1. The protein complex structure prediction protocols

In this study, we only focus on predicting heterodimeric complex structures of intra-species PPIs, for there is no clear way to obtain the interologs of inter-species PPIs. Besides, the prediction of homo-oligomeric complex structure is also not covered by our study, for the MSA of interologs of the homo-oligomeric complex can be built trivially by repeatedly concatenating the MSA of homologous monomers.

The basic complex structure prediction protocol is summarized in Figure 1. As it is shown in the figure, given sequences of the interacting proteins, we first applied JackHMMER[25] to search against UniRef100[26] (2021-04-07) to generate the MSA of homologous proteins for each protein. The UniRef100 database was used for it is the largest sequence database with the compete annotation of species information for each sequence. Then each MSA was filtered by removing sequences with more than 50% gaps, with non-standard amino acids, without species information and with “uncertain”, “uncharacterized” and “putative” in their cluster names, and the kept sequences are grouped according to their species. After that, a simplified phylogeny-based approach was applied to pair sequences belonging to the same species in the two MSAs. Specifically, sequences with the highest similarities to the two interacting proteins were paired to form the (putative) interolog of the target PPI from each species, considering which are more likely to be the orthologs of the two interacting proteins, thus have a higher probability to interact with each other and are also more likely to share the binding mode of the target interacting proteins (see Figure S1). Linkers with each formed of 300 GLYs were then applied to connect the paired sequences to form the MSA of the interologs (phylogeny-MSA), which was inputted to the fine-tuned model 5 (model_5_ptm) of AF2 with the default settings to predict the complex structure of the interacting proteins without using any template information. The fine-tuned model rather the original models of AF2 was used in our study, for which allows us to use pTM to evaluate the global quality of the predicted models. Five fine-tuned models were released in total in AF2. According to the test on a small dataset, the five models achieved very similar performances on protein complex structure prediction, but among them, the model_5_ptm achieved a slightly better performance, thus which was chosen for protein complex structure prediction in our study. In the following sections, this basic protein complex structure prediction protocol is referred to as “AF2 + phylogeny-MSA”. Besides, the genomic distanced-based MSA of interologs produced by the RoseTTAFold protocol (genome-MSA) and the phylogeny-based MSA of interologs produced by the ComplexContact protocol (phylogeny-MSA (all pairs)) were also inputted to the same deep learning model of AF2 to predict protein complex structures respectively for references, which are referred to as “AF2 + genome-MSA” and “AF2 + phylogeny-MSA (all pairs)” in the following sections (see Supplemental Methods S2 and S4 for details).

**Figure 1.**
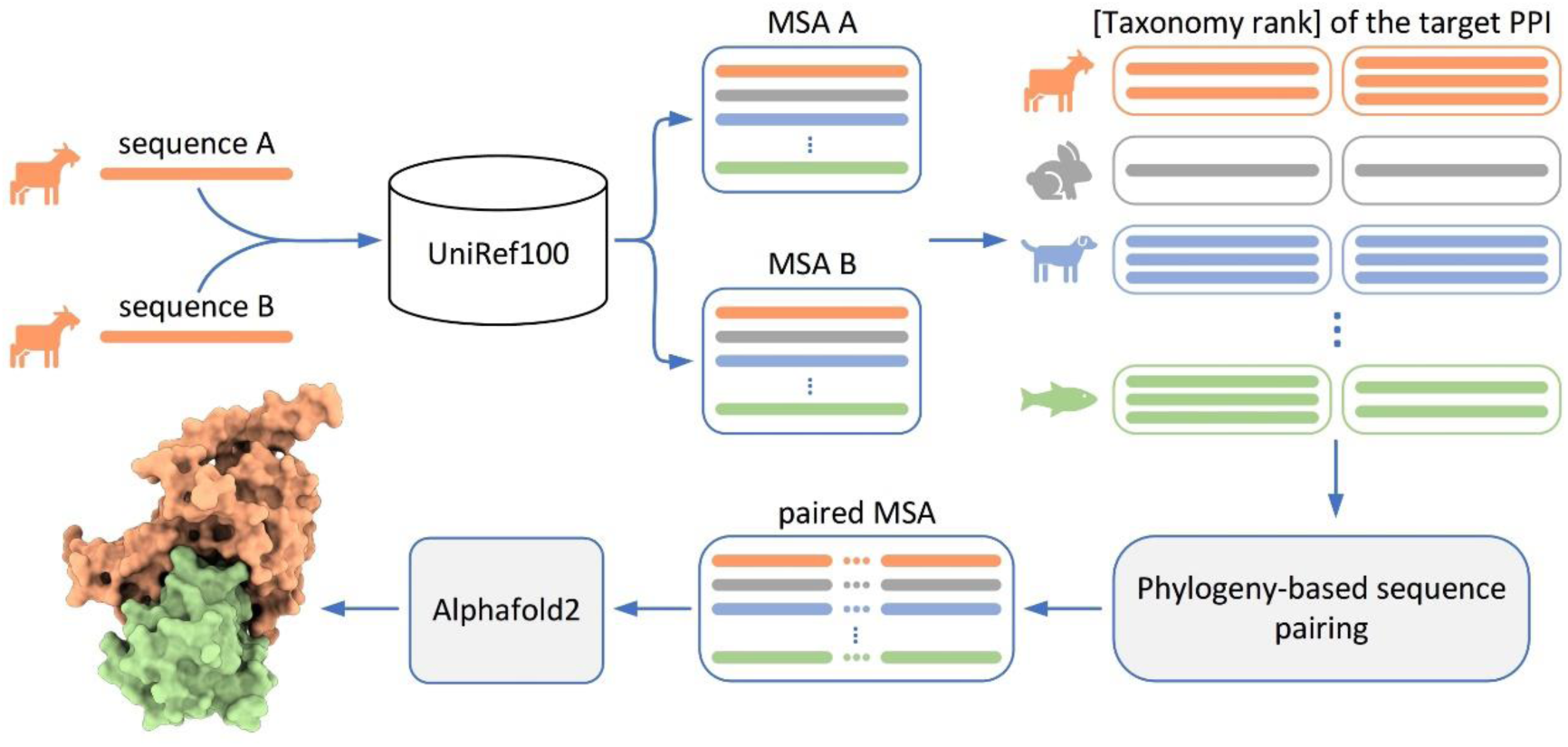
The flowchart of protein complex structure with the application of the phylogeny-based MSA of interologs and AlphaFold2.

Since PPIs are generally not as conserved as monomeric proteins in the evolution process, including sequences of non-conserved protein pairs in the MSA may attenuate the prediction performance. Therefore, we further applied MSAs in which the interologs are from species within different taxonomic ranks (domain, kingdom, phylum, class and order)[27] of the species of the target PPI in the complex structure prediction (See Supplemental Methods S3). Different target PPIs may have different optimal taxonomic ranks in the complex structure prediction. In this study, the protein complex structure prediction with the application of the MSA of interologs from the optimal taxonomic rank is referred to as “AF2 + phylogeny-MSA + rank”. Since the pTM for each generated model can be used to estimate the accuracy of the generated model, we applied pTM to select the predicted complex structure from the multiple generated models on different taxonomic ranks (i.e. the model with the highest pTM was selected as the predicted complex structure), which is referred to as “AF2 + phylogeny-MSA + pTM-rank” in this study. It should be noted that when using pTM for the model selection, we recalculated pTM for each generated model with the function provided in AF2 after removing the GLY linker region in the model.

### 2. The evaluation protocols

The protein complex structure prediction protocols described above were extensively evaluated on non-redundant intra-species PPIs including 107 PPIs in bacteria and 442 PPIs in eukaryotes with experimentally resolved complex structures (see Supplemental Methods S1 for the dataset preparation). For each PPI, we employed DockQ , a continuous quality score in the range [0,1], to assess the quality of the complex structure prediction (see Supplemental Methods S5)[28]. Specifically, DockQ < 0.23 corresponds to an incorrect prediction; 0.23 ≤ DockQ < 0.49 corresponds to an acceptable prediction; 0.49 ≤ DockQ < 0.8 corresponds to a medium quality prediction; and 0.8 ≤ DockQ corresponds to a high quality prediction.

## Results

### 1. Protein complex structure prediction with “AF2 + phylogeny-MSA”

We first applied “AF2 + phylogeny-MSA” to predict complex structures for non-redundant intra-species PPIs including 107 bacterial PPIs and 442 eukaryotic PPIs, and “AF2 + genome-MSA” and “AF2 + phylogeny-MSA (all pairs)” were also applied on the same datasets for reference. In Figure 2, we show the performance comparison between these “AF2 + phylogeny-MSA” and the two reference protocols on protein complex structure prediction. As we can see from Figure 2 that for both the bacterial and the eukaryotic PPIs, “AF2 +phylogeny-MSA” achieved much better performance than “AF2 + genome-MSA”, and slightly better performance than “AF2 + phylogeny-MSA (all pairs)”. Specifically, “AF2 + phylogeny-MSA” was able to successfully predict complex structures for 79.5% of the bacterial PPIs (acceptable quality: 10.3%, medium quality: 39.3%, high quality: 29.9%, mean DockQ: 0.574) and 49.8% of the eukaryotic PPIs (acceptable quality: 10.2%, medium quality: 24.0%, high quality: 15.6%, mean DockQ: 0.355); “AF2 + genome-MSA” only successfully predicted complex structures for 56.1% of the bacterial PPIs (acceptable quality: 5.61%, medium quality: 28.0%, high quality: 22.4%, mean DockQ: 0.426) and 17.6% of the eukaryotic PPIs (acceptable quality: 5.43%, medium quality: 8.59%, high quality: 3.62%, mean DockQ: 0.131); “ AF2 + phylogeny-MSA (all pairs)” successfully predicted complex structures for 78.5% of the bacterial PPIs (acceptable quality: 6.54%, medium quality: 39.3%, high quality: 32.7%; mean DockQ: 0.592) and 47.0% of the eukaryotic PPIs (acceptable quality: 10.2%, medium quality: 22.9%, high quality: 14.0%, mean DockQ: 0.332). Therefore, we can conclude from the result that for both the bacterial PPIs and the eukaryotic PPIs, the simplified phylogeny-based approach is significantly more effective than the genomic distance-based approach in preparing the MSA of interologs to assist protein complex structure prediction, and the inclusion of the lower ranking proteins pairs can even attenuate the protein complex structure prediction performance (see Figure S2 for the statistical tests). It is worth noting that in the work of ComplexContact, a web server for predicting inter-protein residue-residue contacts, the authors claimed that genome-distance approach is more effective in preparing MSA of interologs for PPIs in bacteria, and the phylogeny-based approach is more effective in preparing MSA of interologs for PPIs in eukaryotes[19]. However, it should be noted the bacterial PPI dataset used to evaluate ComplexContact is from Ovchinnikov *et al*.[17], which is constituted by Ecoli PPIs formed by proteins with short genomic distances, but our bacterial PPIs do not have genomic distance restrictions.

**Figure 2.**
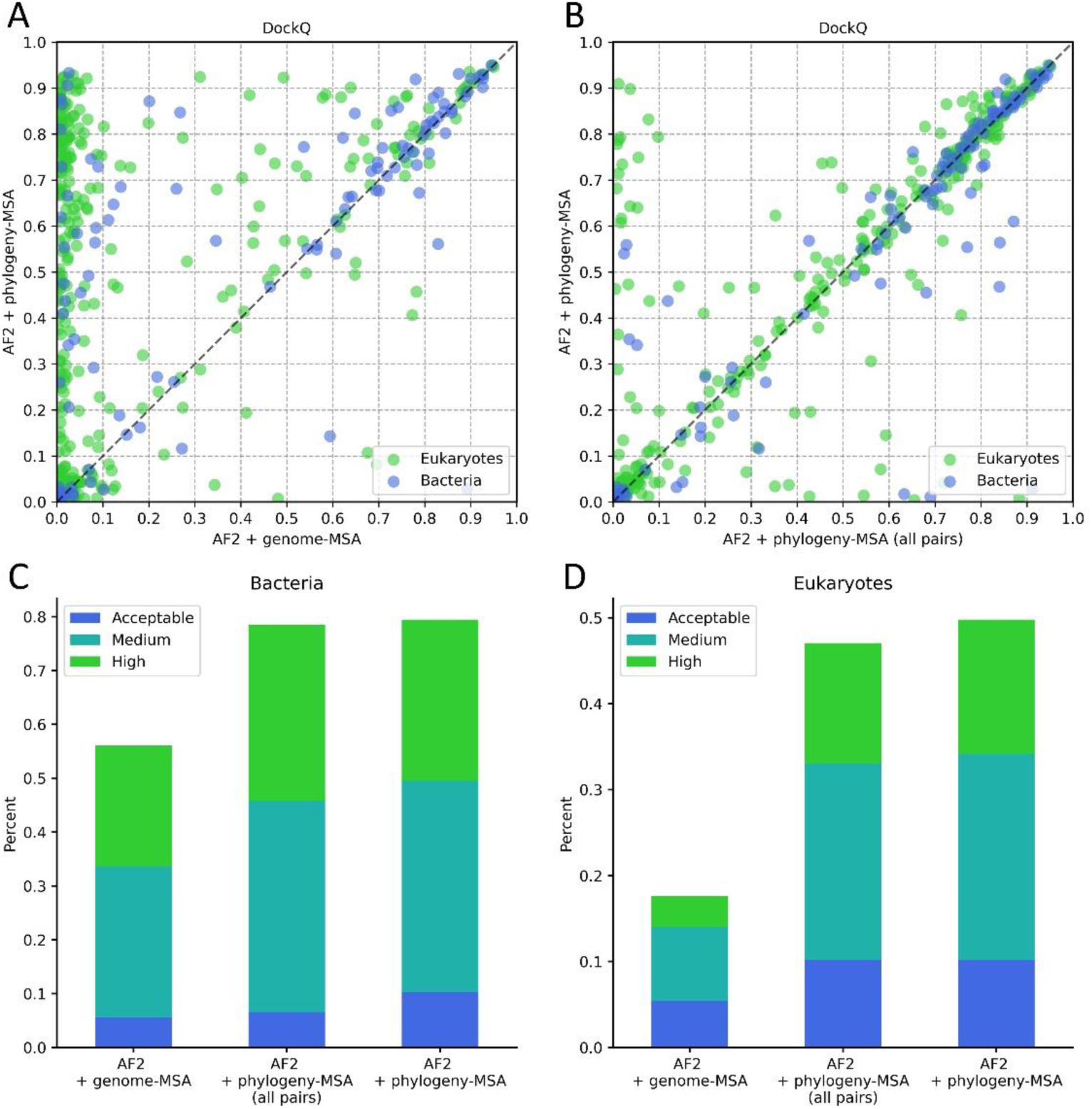
Protein complex structure prediction with the application of the MSA of interologs prepared by different approaches. (A) “AF2 + phylogeny-MSA” versus “AF2 + genome-MSA” for each target PPI. (B) “AF2 + phylogeny-MSA” versus “AF2 + phylogeny-MSA (all pairs)” for each target PPI. (C) Performance comparison between “AF2 + genome-MSA”, “AF2 + phylogeny-MSA (all pairs)” and “AF2 + phylogeny-MSA” on the bacterial PPI dataset. (D) Performance comparison between “AF2 + genome-MSA”, “AF2 + phylogeny-MSA (all pairs)” and “AF2 + phylogeny-MSA” on the eukaryotic PPI dataset.

### 2. Restriction of interologs in the MSA to different taxonomic ranks of the species of the target PPI in protein complex structure prediction

A PPI may not be maintained in another species with very long evolutionary distance, or even though the PPI is maintained, the complex structure of the interolog can be different, for PPIs are often not as conserved as monomeric proteins in the evolutionary process[23, 24]. Including sequences of non-conserved protein pairs in the MSA may attenuate the prediction performance. Therefore, we further applied MSAs in which the interologs are from species within different taxonomic ranks (domain, kingdom, phylum, class and order) of the species of the target PPI in the complex structure prediction. In Figure 3A-E, we show the performance comparison between the protein complex structure prediction with application of MSA of interologs from each taxonomic rank (“AF2 + phylogeny-MSA (domain/kingdom/phylum/class/order)”) and from the whole UniRef100 database (“AF2 + phylogeny-MSA”). As we can see from Figure 3A-E that higher prediction accuracies for a significant number of PPIs can be achieved when restricting interologs in the MSA to each taxonomic rank. Different target PPIs may have different optimal taxonomic ranks for protein complex structure prediction. Of course, for some PPIs, the best prediction performance was achieved when there was no taxonomic rank restriction. In our study, “life” was used to represent all species in the UniRef100 database, which can be considered as the highest taxonomic rank. If the optimal taxonomic rank can be chosen for each target PPI in protein complex structure prediction, higher quality predictions can be obtained for a large number of PPIs (see Figure 3E). Specifically, with the application of the MSA of interologs from the optimal taxonomic rank for each target in protein complex structure prediction (“AF2 + phylogeny-MSA + rank”), the success rates for protein complex structure prediction can be increased to 87.9% for the bacterial PPIs (acceptable quality: 10.3%, medium quality: 38.3%, high quality: 39.3%, mean DockQ:0.652) and 56.3% for the eukaryotic PPIs (acceptable quality: 10.4%, medium quality: 25.8%, high quality: 20.1%, mean DockQ: 0.406) (see Figure 4A-B and also see Figure S3 for the statistical tests). An example of the improvement is shown in Figure 4C-D. As we can see from Figure 4C-D, with the application of MSA of interologs from the whole UniRef100 database as the input to AF2, an incorrect model (DockQ=0.03) for the PPI in PDB 3NK8 was generated in the protein complex structure prediction. However, when we restricted interologs in the MSA to the class of the target PPI, a high quality prediction (DockQ=0.89) was achieved.

**Figure 3.**
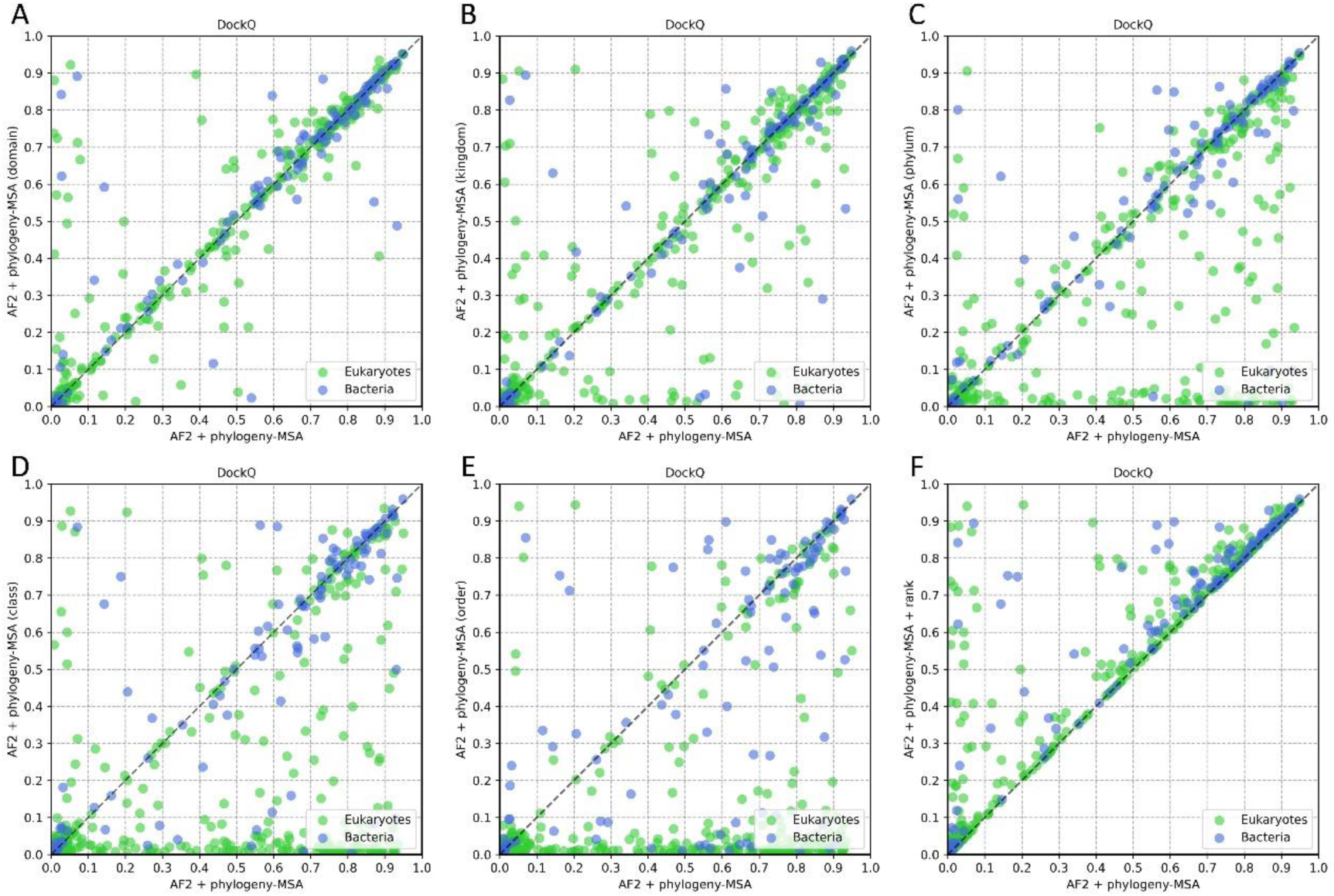
Protein complex structure prediction with the restriction of interologs in the MSA to different taxonomic ranks of species of the target PPI. (A) “AF2 + phylogeny-MSA (domain)” versus “AF2 + phylogeny-MSA” for each target PPI. (B) “AF2 + phylogeny-MSA (kingdom)” versus “AF2 + phylogeny-MSA” for each target PPI. (C) “AF2 + phylogeny- MSA (phylum)” versus “AF2 + phylogeny-MSA” for each target PPI. (D) “AF2 + phylogeny-MSA (class)” versus “AF2 + phylogeny-MSA” for each target PPI. (E) “AF2 + phylogeny-MSA (order)” versus “AF2 + phylogeny-MSA” for each target PPI. (F) “AF2 + phylogeny-MSA + rank” versus “AF2 + phylogeny-MSA” for each target PPI.

**Figure 4.**
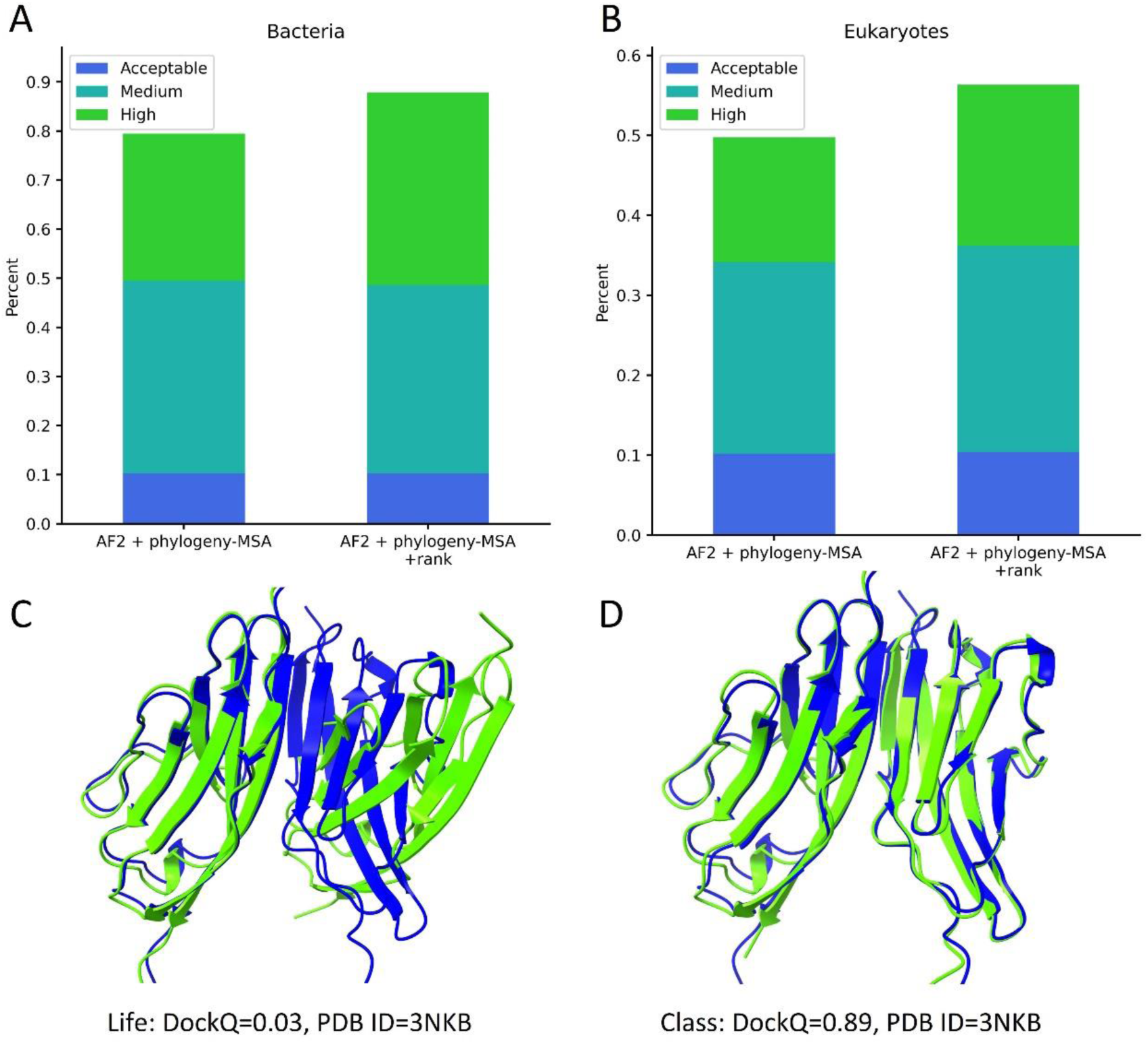
Protein complex structure prediction with the restriction of interologs in the MSA to the optimal taxonomic rank of the species of each target PPI. (A) Performance comparison between “AF2 + phylogeny-MSA” and “AF2 + phylogeny-MSA + rank” on the bacterial PPIs. (B) Performance comparison between “AF2 + phylogeny-MSA” and “AF2 + phylogeny-MSA + rank” on the eukaryotic PPIs. (C) The quality of the predicted model for PDB 3KNB when the MSA of interologs was not restricted to a specific taxonomic rank of the species of the target PPI. (D) The quality of the predicted model for PDB 3KNB when the MSA of interologs was restricted to the class of the target PPI.

### 3. Application of pTM to select taxonomic ranks for protein complex structure prediction

The application of the fine-tuned model of AF2 allows us to use pTM to evaluate the quality of each predicted model. Apart from the pTM, AF2 also outputs pLDDT (predicted IDDT-Cα, ranging from 0-100) to measure the per-residue local prediction confidence along the protein chain, and the mean value of pLDDT is used as the default by AF2 to evaluate the overall prediction accuracy of each predicted model. For monomeric protein structure prediction, both the pTM and the mean pLDDT can be used to evaluate the prediction quality. However, pLDDT is not appropriate to evaluate the quality of the predicted complex structures, since pLDDT mainly measures the local prediction confidence and cannot reflect the global accuracy of the predicted complex structures. For example, even when the monomeric structures of the two interacting proteins are accurately predicted, the predicted binding modes between them can be totally wrong. In this case, the mean pLDDT is still very high, for the pLDDT values along the two interacting chains are high. We further confirmed this on the generated models (also using MSA of interologs from different taxonomic ranks) for the 31 PPIs in our PPI dataset which were deposited in PDB after the training of AF2 (April 30, 2018). We found that even when the mean pLDDT (excluding the linker region) is higher than 80, the predicted complex structure still has a high chance (43%) to be with incorrect conformation (DockQ<0.23), however, when pTM>0.8 (the pTM was also recalculated after removing the linkers), most of the predicted complex structures (95%) are at least with acceptable qualities (DockQ≥0.23) (see Figure S4). Actually, the performance of mean pLDDT would be even worse if models of PPIs before the training set of AF2 were included, for the interacting proteins of these PPIs were included in the AF2 training, thus the pLDDT values would be consistently high across the monomeric chains.

Therefore, pTM was applied to select the taxonomic ranks (including the taxonomic rank of “life”) for protein complex structure prediction in this study. Specifically, we predicted protein complex structure for each PPI with the application of MSAs of interologs from the six different taxonomic ranks (life/domain/kingdom/phylum/class/order) of the target PPI respectively, thus six different predicted models can be generated for each PPI. Then we recalculated the pTM for each predicted model with the function provided in AF2 after the removal of the GLY linker, and the model with the highest recalculated pTM was considered as the final predicted model for the corresponding PPI (“AF2 + phylogeny-MSA + pTM-rank”). In Figure 5, we show the relationship between the recalculated pTM and the DockQ of the predicted models. As we can see from the figure, the model with the higher pTM tends to have higher DockQ (i.e. higher quality)), especially when the pTM is high (e.g. pTM>0.8), which further supports the application of pTM for the model selection. In Figure 6, we show the performance of “AF2 + phylogeny-MSA +pTM-rank” in protein complex structure prediction, which is also compared with “AF2 + phylogeny-MSA” and “AF2 + phylogeny-MSA + rank”. As we can see from the figure, for only a relatively small number of PPIs, the predicted models by “AF2 + phylogeny-MSA + pTM-rank” show significantly lower qualities than those by “AF2 + phylogeny-MSA +rank”, but for a quite a significant number of PPIs, the predicted models by “AF2 + phylogeny-MSA + pTM-rank” show significantly higher qualities (see Figure 5A-B and also see Figure S5 for the statistical tests). Specifically, with the application of interologs from the pTM-selected taxonomic ranks in the MSA generation, we can successfully predict protein complex structures for 84.2% of the bacterial PPIs (acceptable quality: 11.2%, medium quality: 38.3%, high quality: 34.6%, mean DockQ: 0.615) and for 54.1% of the eukaryotic PPIs (acceptable quality: 10.4%, medium quality: 26.7%, high quality: 17.0%, mean DockQ: 0.382) ( see Figure 5C-D).

**Figure 5.**
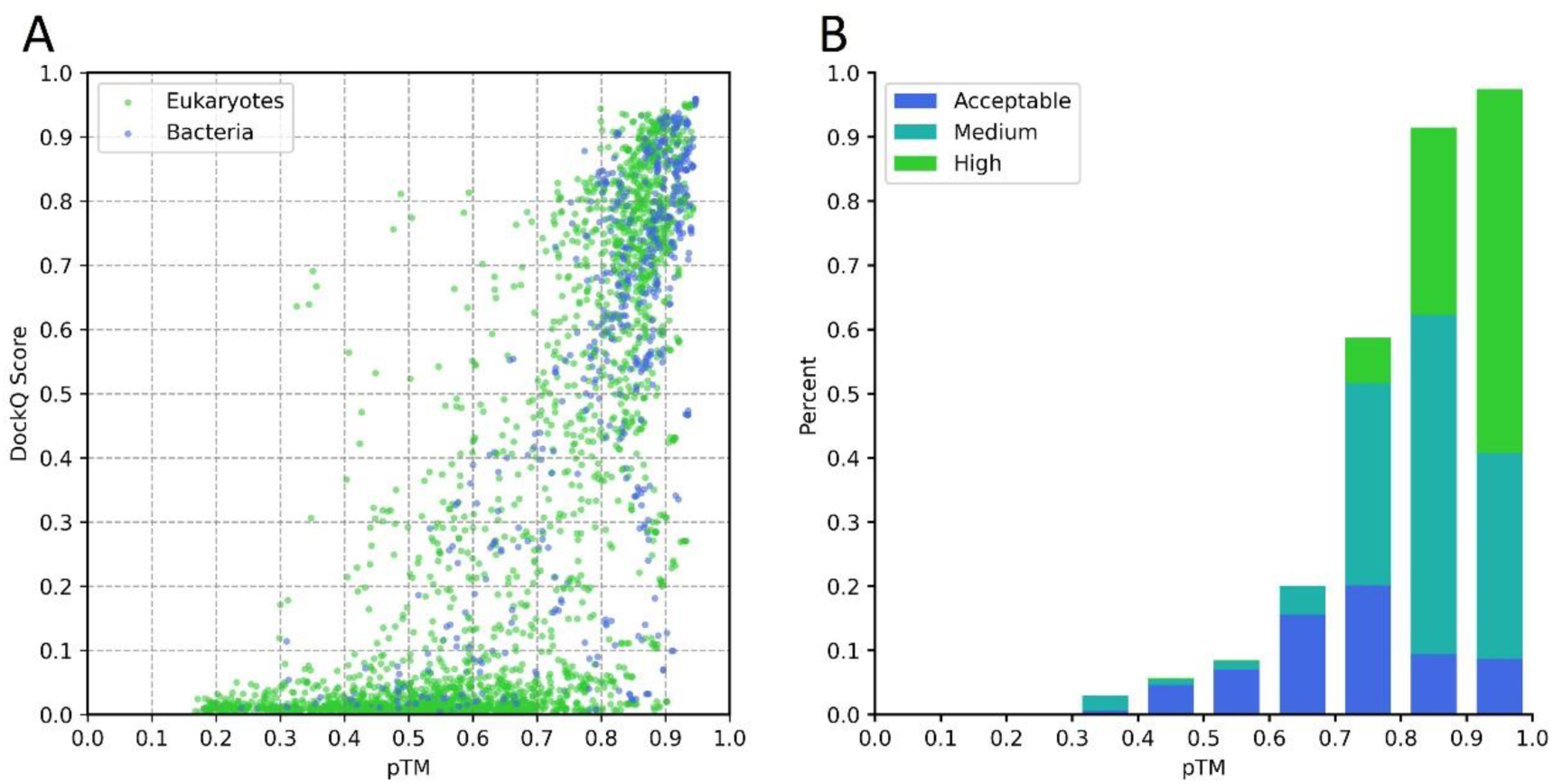
The relationship between the pTM and the qualities of the predicted models. (A) DockQ versus pTM for each predicted model, all the generated models using the MSAs of interologs from different taxonomic ranks of the species of the target PPIs are included in the figure. (B) The proportions of the models with acceptable quality, medium quality and high quality when the pTM values are within different intervals.

**Figure 6.**
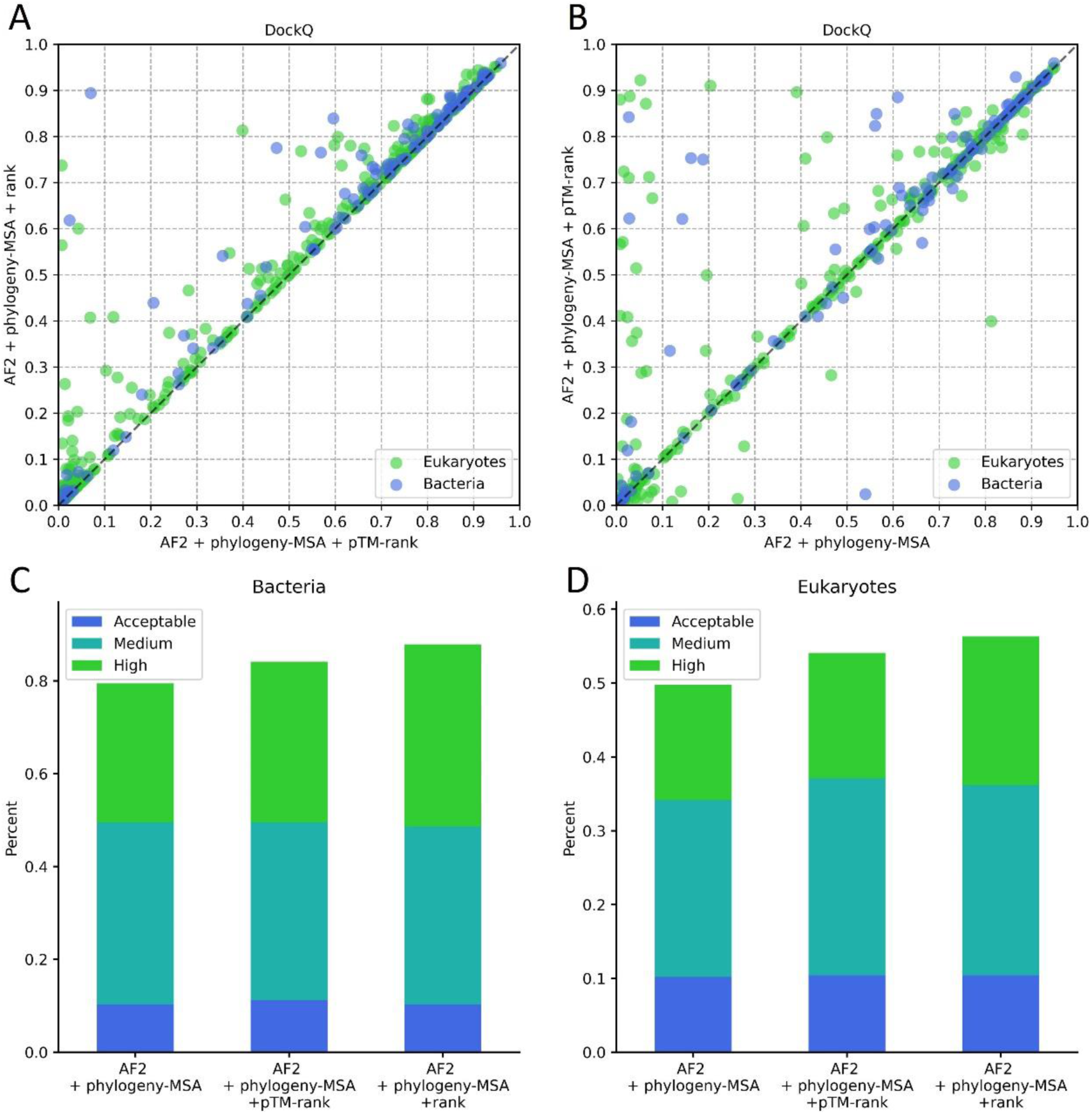
Protein complex structure prediction with the restriction of interologs in the MSA to the pTM-selected taxonomic rank of the species of each target PPI. (A) “AF2 + phylogeny-MSA + pTM-rank” versus “AF2 + phylogeny-MSA + rank” for each target PPI. (B) “AF2 + phylogeny-MSA + pTM-rank” versus “AF2 + phylogeny-MSA” for each target PPI. (C) Performance comparison between “AF2 + phylogeny-MSA”, “AF2 + phylogeny- MSA + pTM-rank” and “AF2 + phylogeny-MSA + rank” on the bacterial PPI dataset. (D) Performance comparison between “AF2 + phylogeny-MSA”, “AF2 + phylogeny-MSA + pTM-rank” and “AF2 + phylogeny-MSA + rank” on the eukaryotic PPI dataset.

### 4. Evaluation of protein complex structure prediction protocols on PPIs deposited after the AF2 training set

Since AF2 was trained on structures of protein monomers deposited in PDB before April 30, 2018, only structures of protein monomers in each PPI have the possibility to be included in the training of AF2, but none of the complex structures would be included in the model training[13]. Besides, all the predictions were made by inputting the MSA of interologs to the deep learning model of AF2 without using any template information. Therefore, it is unlikely that the protein complex structure prediction was made by trivially recognizing the existing templates in PDB or due to the bias of the deep learning model to the target PPIs. However, the inclusion of the interacting protein monomers in the training of AF2 can also cause the overestimation of the model performance in protein complex structure prediction. To provide a more objective evaluation of these protein complex structure prediction protocols, we further evaluated their performances only on the 31 PPIs in our dataset (28 eukaryotic PPIs and 3 bacterial PPIs) with their complex structures deposited in PDB after April 30, 2018. In Figure 7, we show the performance comparison between different protein complex structure prediction protocols on the 31 PPIs (see Figure S6 for the statistical tests). As we can see from the figure that the performance comparison between these protocols on the 31 PPIs is consistent with those on the full dataset. Specifically, the mean DockQ of the 31 predicted models by “AF2 + phylogeny-MSA” is 0.295 (acceptable quality: 16.1%, medium quality: 19.4%, high quality: 9.68%), which is significant higher than the performance of “AF2 + genome-MSA” (mean DockQ: 0.05, acceptable quality: 3.22%, medium quality: 3.22%, high quality: 0%) and the performance of “AF2 + phylogeny-MSA (all pairs)” (mean DockQ: 0.239, acceptable quality: 16.1%, medium quality: 3.22%, high quality: 16.1%) ; by further restricting interologs in the MSA to the optimal taxonomic ranks (“AF2 + phylogeny-MSA + rank”), the mean DockQ of the predicted models can be increased to 0.345 (acceptable quality: 19.4%, medium quality: 12.9%, high quality: 19.4%); with the application the pTM-selected taxonomic ranks for protein complex structure prediction (“AF2 + phylogeny-MSA + pTM-rank”), the mean DockQ of the predicted models is 0.324 (acceptable quality: 22.6%, medium quality: 12.9%, high quality: 16.1%), which is only slightly lower than the ideal case (i.e. “AF2 + phylogeny-MSA + rank”). The performances of these protein complex structure prediction protocols on the more recent 31 PPIs (28 of which are eukaryotic PPIs) are similar to the performances on the full eukaryotic PPI dataset. Therefore, arguably we can say that it has very little overestimation of the performances of these protein complex structure prediction protocols when they were evaluated on the full PPI dataset.

**Figure 7.**
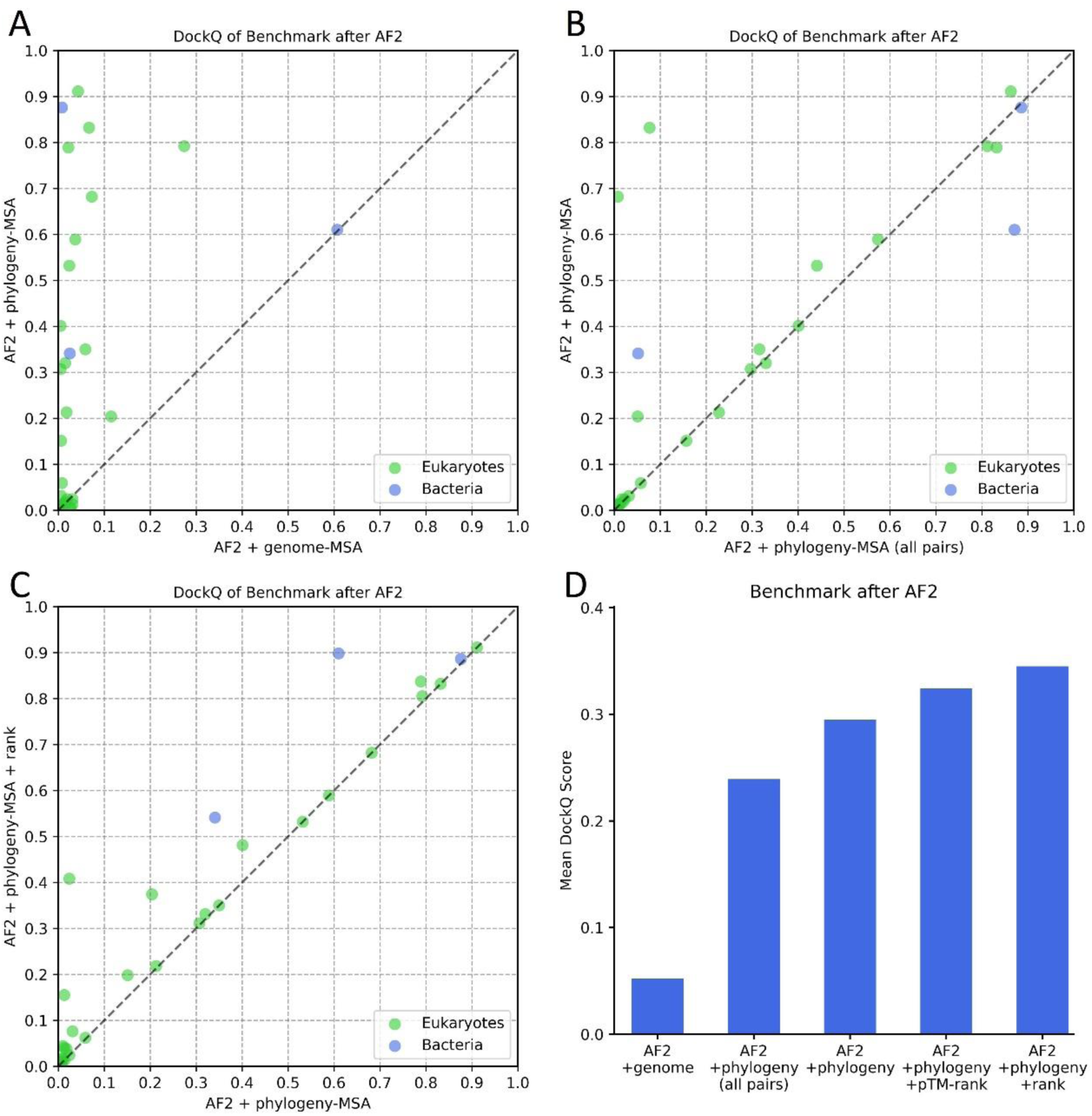
Protein complex structure prediction on PPIs after the training of AF2. (A) “AF2 + genome-MSA” versus “AF2 + phylogeny-MSA” for each target PPI. (B) “AF2 + phylogeny-MSA” versus “AF2 + phylogeny-MSA (all pairs)” for each target PPI. (C) “AF2 + phylogeny-MSA + rank” versus “AF2 + phylogeny-MSA” for each target PPI. (D) The mean DockQ scores of the predicted complex structures with the application of different protein complex structure prediction protocols.

### 5. The successful cases versus the failed cases

As it was shown above, even when the optimal taxonomic rank was chosen in the MSA generation, our protocol (i.e. “AF2 + phylogeny-MSA + rank”) still failed to predict protein complex structures for 12.1% of the bacterial PPIs and 43.7% of the eukaryotic PPIs. To explore the possible reasons accounting for the failures in the protein complex structure prediction, both the bacterial PPIs and the eukaryotic PPIs were grouped into two subsets respectively based on whether the complex structure of the PPI can be successfully predicted (DockQ ≥ 0.23). We found that the failed cases of both the bacterial PPIs and eukaryotic PPIs tend to have fewer effective sequences in their MSAs. As it is shown in Figure 8A, for the bacterial PPIs, the median value of the normalized effective sequence number of the MSA (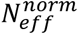) of the failed cases is only 1.08 (see Supplemental Methods S6 for the calculation of 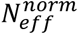), which is dramatically lower than the median value of 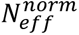 of the successful cases (median 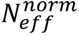 = 12.85) . For the eukaryotic PPIs, although the median value of 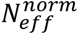 of the failed cases is not as low as that of the bacterial PPIs (median 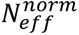 = 6.48), which is still significantly lower than that of the successful cases (median 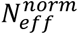=10.87). Besides, we also found that the failed cases of the eukaryotic PPIs tend to have significantly smaller inter-protein interfaces, but the interfaces of the failed cases of the bacterial PPIs are with similar sizes to the interfaces of the successful cases. As it shown in Figure 8B, the median number of inter-protein contacts (a contact is defined if the heavy atom distance of two residues is within 5Å) of the failed cases of the eukaryotic PPIs is 34, which is significantly lower than that of the successful cases (median number is 74), and for the bacterial PPIs, the median numbers of inter-protein contacts of the successful cases and failed cases are 89 and 99 respectively. Obviously, for the failed cases of bacterial PPIs, the lack of effective sequences in the MSA is the main reason for the failures in the protein complex structure prediction. For the failed cases of the eukaryotic PPIs, apart from the lack of effective sequences in their MSAs, we doubt the relatively smaller PPI interfaces are also responsible for the failures. Since the deep learning model of AF2 was trained on protein monomers, which may not perform well on modelling transient PPIs with small interfaces. When this manuscript is under preparation, DeepMind also released their deep learning models trained on protein complex structures[29], it would be interesting to see whether our failed cases can be correctly predicted by inputting our MSAs to the new deep learning models. Besides, since eukaryotic proteins generally contain much more paralogs than prokaryotic proteins, our phylogeny-based sequence pairing protocol may not work well for some eukaryotic PPIs for it is more challenging to select orthologs from homologous sequences. Therefore, we doubt that mis-paired proteins with no interactions may be included in forming the MSA of interologs for some eukaryotic PPIs, which can also be the reason that the protein complex structure prediction was failed for some PPIs in which the MSAs contain a large number of paired sequences. It would also be interesting to see whether the prediction performance can be further improved if more rigorous and complex protocols are used for ortholog prediction. Besides, in our study, only the sequence with the highest similarity to each target interacting protein in each specific species (i.e. the putative ortholog) was included in forming the MSA of protein-protein interologs, appropriately including the lower ranking sequences (i.e. the putative paralogs) in the MSA generation may also improve the protein complex structure prediction. However, in our study, we found that including the lower ranking sequences (i.e. the putative paralogs) with the protocol of ComplexContact does not help the protein complex structure prediction. Therefore, developing more effective protocols to pair the putative paralogs of interacting proteins can also be a future direction to improve the protein complex structure prediction performance.

**Figure 8.**
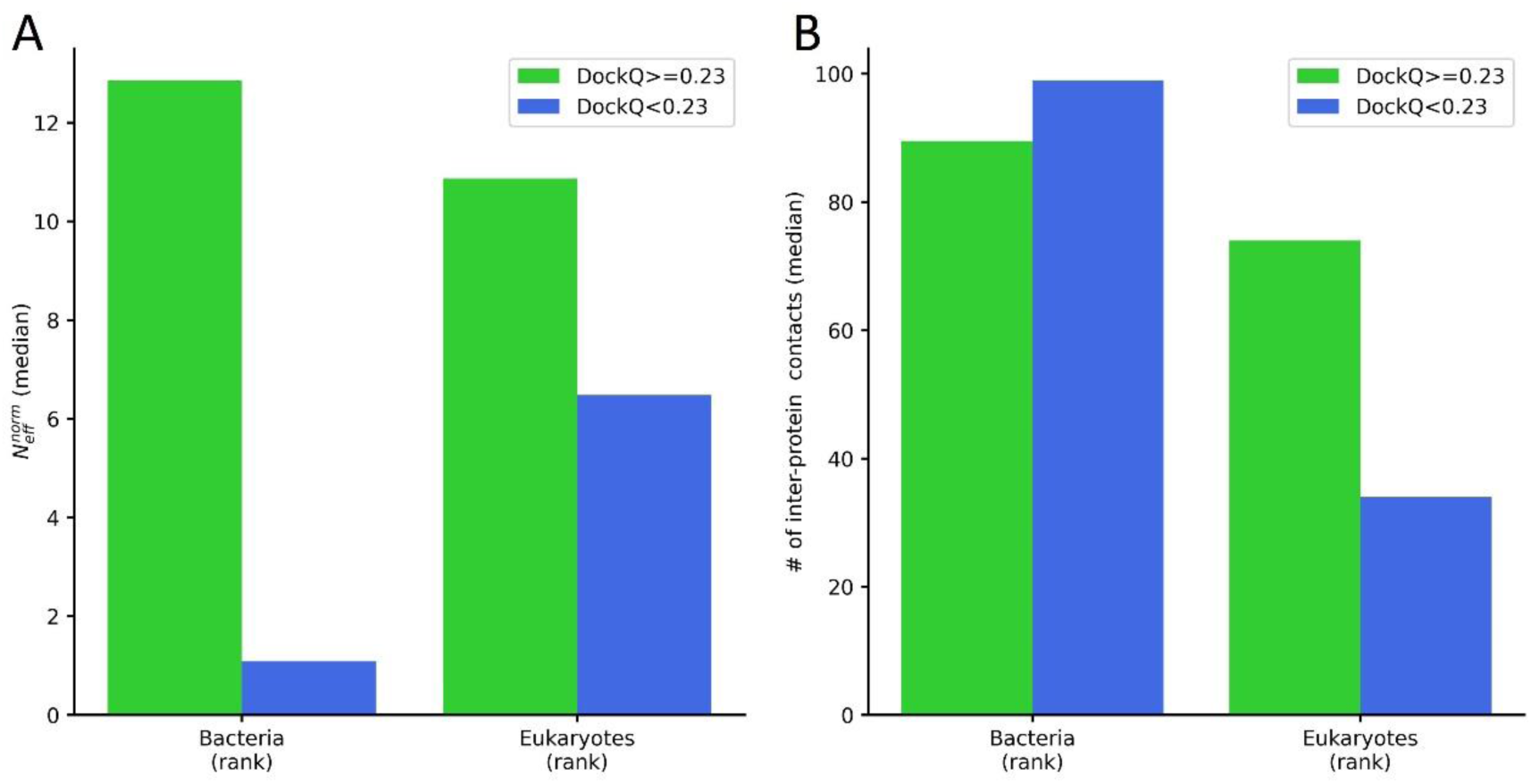
The successful cases versus the failed cases in protein complex structure prediction with “AF2 + phylogeny-MSA + rank”. (A) The comparison of the median values of 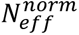 for the successful cases and the failed cases in the bacterial PPI dataset and eukaryotic PPI dataset respectively. (B) The comparison of the median value of the number of inter-protein contacts for the successful cases and the failed cases in the bacterial PPI dataset and eukaryotic PPI dataset respectively.

## Conclusion and Discussions

The deep learning architecture of AF2 is expected to be able to predict complex structures of PPIs, as long as the MSA of interologs of the target PPI can be provided as the input. However, preparing the MSA of interologs is not trivial due to the existence of paralogs. In this study, a simplified phylogeny approach was employed to generate the MSA of interologs for intra-species PPIs, in which the sequences with the highest similarity to the interacting proteins in each specific species are paired to form the MSA of interologs. Benchmarked on large dataset of bacterial and eukaryotic intra-species PPIs with experimentally resolved complex structures, we show that the complex structures of most of the bacterial PPIs and more than half of the eukaryotic PPIs can be successfully predicted by inputting the MSA of interologs prepared by this simplified phylogeny-based approach into the deep learning model of AF2. We further show that the prediction performance can be significantly improved if we only include interologs from species belonging to a specific taxonomic rank of the species of the target PPI in the MSA generation, and the pTM of the predicted models can be used to select the optimal taxonomic ranks for protein complex structure prediction. Future works on improving the protein complex structure prediction performance include developing more effective protocols to generate the MSA of interologs and employing more accurate deep learning models trained specifically from protein complex structures for protein complex structure prediction. Besides, protein complex structure prediction for inter-species PPIs was not covered by this study. For inter-species PPIs, preparing the MSA of interologs is challenging, since the intra-species wised pairing mechanism may not work well for inter-species PPIs. In this case, we may not be able to use AF2 to directly predict the complex structure of the PPI. However, AF2 can still be used to determine the monomeric structure of each interacting protein, and then traditional protein complex structure prediction methods like protein-protein docking can be employed to predict the protein complex structure. Besides, a recent study shows that protein monomeric structures can be predicted with high accuracy by deep learning irrespective of co-evolution information [30]. We may also be able to develop deep learning models for protein complex structure prediction without leveraging inter-protein coevolutionary information (i.e. without using MSA of interologs).

## Key points

- Protein complex structures are predicted by inputting MSAs of interologs to AlphaFold2.
- MSA of interologs prepared by a simplified phylogeny approach outperforms two existing approaches
- Restricting interologs to certain taxonomic ranks improves prediction performance.
- The predicted TM-scores can be used to select the optimal taxonomic ranks for protein complex structure prediction.

## Code Availability

The code for generating the MSA of interologs for a given PPI can be obtained from https://github.com/ChengfeiYan/PPI_MSA-taxonomy_rank.

## Funding

The work was supported by the National Natural Science Foundation of China (32101001) and new faculty startup grant (3004012167) of Huazhong University of Science and Technology.

**Yunda Si** is a PhD student in the School of Physics at Huazhong University of Science and Technology. His research interests include protein structure prediction, protein-protein interaction prediction and deep learning.

**Chengfei Yan** is an associate professor in the School of Physics at Huazhong University of Science and Technology. His research interests include molecular docking, protein-protein interaction prediction and biological data mining.

