## Supplemental Figures and text for "Protein Complex Structure Prediction Powered by Multiple Sequence Alignments of Interologs from Multiple Taxonomic Ranks and AlphaFold2"

### Supplementary

### Supplemental Methods

#### S1. Preparing the non-redundant PPI dataset

##### 1.1 Collecting protein complex structures

To prepare the PPI dataset used in this study, protein complex structures were first collected from the Protein Data Bank[1]. Specifically, the criteria including 1) Experimental Method equals to X-RAY DIFFRACTION; 2) Data Collection Resolution $\leq$2.5Å; 3) Entry Polymer Types equals to Protein (only); 4) Entry Polymer Composition equals to heteromeric protein; 5) Total Number of Polymer Instances $\geq 2$; 6) SCOP classification belongs to [(1) All beta proteins, (2) All alpha proteins, (3) Alpha and beta proteins (a+b), (4) Alpha and beta proteins (a/b), (5) Membrane and cell surface proteins and peptides] were used to filter the experimental structures deposited in PDB before July 7, 2021. In total, 9509 protein complex structures were obtained from the initial filter.

##### 1.2 Extracting complex structures of PPIs from protein complex structures

Then complex structures of PPIs were then extracted from the protein complex structures collected above. Specifically, protein complex structures with more than one assembly were removed in our study, for complex structures of PPIs cannot be uniquely determined from the corresponding protein complex structures. For protein complex structures with only two protein chains, the protein complex structure directly corresponds to the complex structure of the two interacting chains. For protein complex structures with more than two chains, the protein complex structure was split into multiple structures in accordance with different combinations of the chains. Then PPIs which satisfied any of the following conditions: 1) the two interacting proteins belong to different species; 2) the PPI belongs to a species without clear taxonomic rank information; 3) either of the interacting protein chains is shorter than 50 or longer than 700; 4) the sequence identity of the two interacting proteins are higher than 40%; 5) either of the two proteins have more than 20% missing residues were removed from the PPI dataset; 6) there are more than 10 contact sites in the two chains (a contact is defined if the heavy atom distance of two residues is within 6Å). After the removal, 2895 complex structures of PPIs can be obtained.

**1.3 Removing redundant PPIs**

Finally, redundant PPIs were removed from the PPI dataset obtained above. Specifically, 5790 proteins in the 2895 PPIs were clustered with CD-HIT[2] using 40% as the sequence identity threshold (-c 0.4 -n 0.2 -g 1), from which 640 clusters were obtained. The obtained clusters were then used to label each PPI. The PPIs belonging to the same cluster pair (note: cluster pair AB and cluster pair BA were considered as the same pair) were considered as redundant PPIs, for which only the PPI with largest number of inter-protein contacts (a contact is defined if the heavy atom distance of two residues is within 6Å) was kept. In total 563 non-redundant PPIs were kept. Among them, 9 are archaea PPIs; 5 are viruses PPIs; 107 are bacterial PPIs; and 442 are eukaryotic PPIs. The PPIs belonging to the two major domains including bacteria and eukaryotes were used to evaluate our protein complex structure prediction protocols in this study.

**S2. Preparing the MSA of interologs**

**2.1 Generating the MSA of homologous proteins for each interacting protein**

To prepare the MSA of interologs for a given PPI, the MSA of homologous proteins for each interacting protein needs to be generated first. Specifically, JackHMMER[3] was applied to search against the UniRef100 database[4] (released on 2021-04-07) with ( --noali --incT L/2 ), where L is the protein length. Then the MSAs for the two interacting proteins were paired to form the MSA of interologs for the PPI with a simplified phylogeny-based approach.

##### 2.2 Pairing the MSAs with the simplified phylogeny-based approach

Before paring the MSAs, we first removed sequences with more than 50% gaps, with non-standard amino acids, without species information and with "uncertain", "uncharacterized" and "putative" in their cluster names in each MSA. After the filter, sequences in each MSA were grouped according to their species, and the sequences of each specific species were sorted descendingly according to their sequence similarities ($\frac{\#Idnetical\ positions}{\#Aligned\ positions}$) to the target sequence. The top ranked sequences within each specific species in the two MSAs were considered as the putative orthologs of the target interacting proteins, which were paired to form the interolog of the target PPI. Finally, linkers with each formed of 300 GLYs were applied to connect the paired sequences to form the MSA of protein-protein interologs, which can be directly inputted to AF2 to predict the complex structure of the target PPI. Apart from the simplified phylogeny-based approach, the phylogeny-based approach of ComplexContact[5] and the genomic distance-based approach of RoseTTAFold[6] were also used as the reference approaches to prepare the MSA of protein-protein interologs.

##### 2.3 Pairing the MSAs with the phylogeny-based approach of ComplexContact

The major difference between the phylogeny-based approach of ComplexContact and our simplified phylogeny-based approach is that in our simplified phylogeny-based approach, only the top ranked sequences within each specific species were paired, however, in the phylogeny-based approach of ComplexContact, the sequences with the same rankings within each specific species were paired (i.e. the lower ranking protein pairs were included in generating the MSA of protein-protein interologs). Since ComplexContact does not release its tool to prepare the MSA of interologs, we implemented this approach by ourselves. Specifically, we employed the same procedures as our simplified phylogeny approaches in generating the MSA for each interacting protein and in filtering homologous sequences. However, in the sequence pairing stage, we included the lower ranking proteins pairs. Still, linkers with each formed by 300 GLYs were used to connect the paired sequences to form the MSA of protein-protein interologs.

**2.4 Pairing the MSAs with the genomic distance-based approach of RoseTTAFold**

The protocol provided by the RoseTTAFold website (https://github.com/RosettaCommons/RoseTTAFold/tree/main/example/complex_modeling) was exactly used to pair the MSAs with the genomic distance-based approach. Specifically, the “make_joint_MSA_bacterial.py” script provided by RoseTTAFold was first applied to pair sequences in the two MSAs according to their UniProt IDs, and then hhfilter was applied to filter the paired alignment (-id 90 -cov 75). To make the paired MSA compatible with AF2 for protein complex structure prediction, linkers with each formed by 300 GLYS were inserted to connect the paired sequences to form the MSA of interologs as well.

**S3. Restricting interologs in the MSA to different taxonomic ranks of the species of the target PPI**

The species and taxonomic ranks information of the proteins were obtained from the NCBI taxonomy database (https://ftp.ncbi.nlm.nih.gov/pub/taxonomy/new_taxdump/)[7]. To restrict interologs in the MSA to a specific taxonomic rank (domain, kingdom, phylum, class and order) of the target PPI, only the sequences belonging to the corresponding taxonomic rank of the species of the target PPI were included in the MSA generation.

**S4. Predicting protein complex structures**

For a given PPI, the MSA of interologs for the PPI was inputted the fine-tuned model 5 (model_5_ptm) of AF2[8] to predict the complex structure of the PPI without using any template information. The fine-tuned model was chosen for we would like to use the pTM to evaluate the predicted models. Five fine-tuned models were released in total in AF2. According to a test on a small dataset, the five models achieved very similar performances on protein complex structure prediction, but among them, the model_5_ptm achieved a slightly better performance, thus which was chosen for protein complex structure prediction in our study. To use our own MSA of interologs rather than the MSA obtained by the search engine of AF2 in protein complex structure prediction, we edited the code of AF2 to replace the default MSA search engine with our own MSA of interologs generation protocols. To evaluate the quality of the predicted complex structure, the pTM of predicted complex structure was recalculated with the function provided in AF2 after the removal of the GLY linker in the predicted complex structure.

#### S5. Calculating DockQ for the predicted complex structure

The DockQ of the predicted complex structures were calculated with the DockQ software ( https://github.com/bjornwallner/DockQ)[9].

#### S6. Calculating $N_{eff}^{norm}$ for the MSA of interologs

For a given MSA of interologs, the GLY linkers in the MSA were first removed. Then the $N_{eff}^{norm}$ for the MSA of interologs was calculated according to:

$$N_{eff}^{norm}=\frac{1}{\sqrt{L}}\sum_{n=1}^{N}\frac{1}{m_n} \quad (1).$$

Where L is the number of columns of the MSA after the removal of the GLY linkers, $\frac{1}{m_n}$ is the weight the n-th interolog with $m_n$ being the number of interologs in the MSA which have a sequence identity higher than 80% to the n-th inerologs; and N is the total number of interologs in the MSA.

#### Supplemental Figures

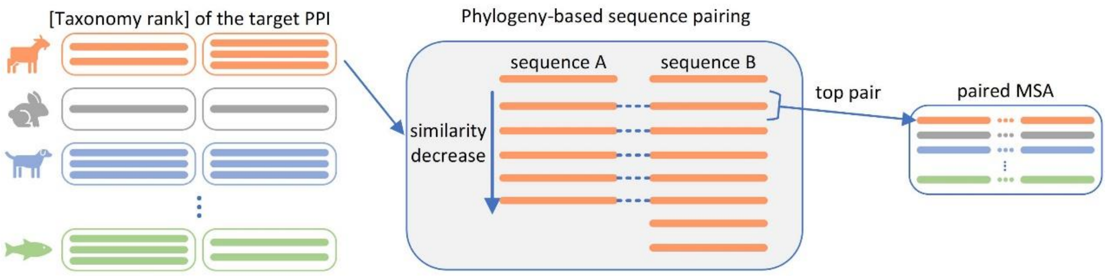


Figure S1. The simplified phylogeny-based approach for MSA of interologs generation.

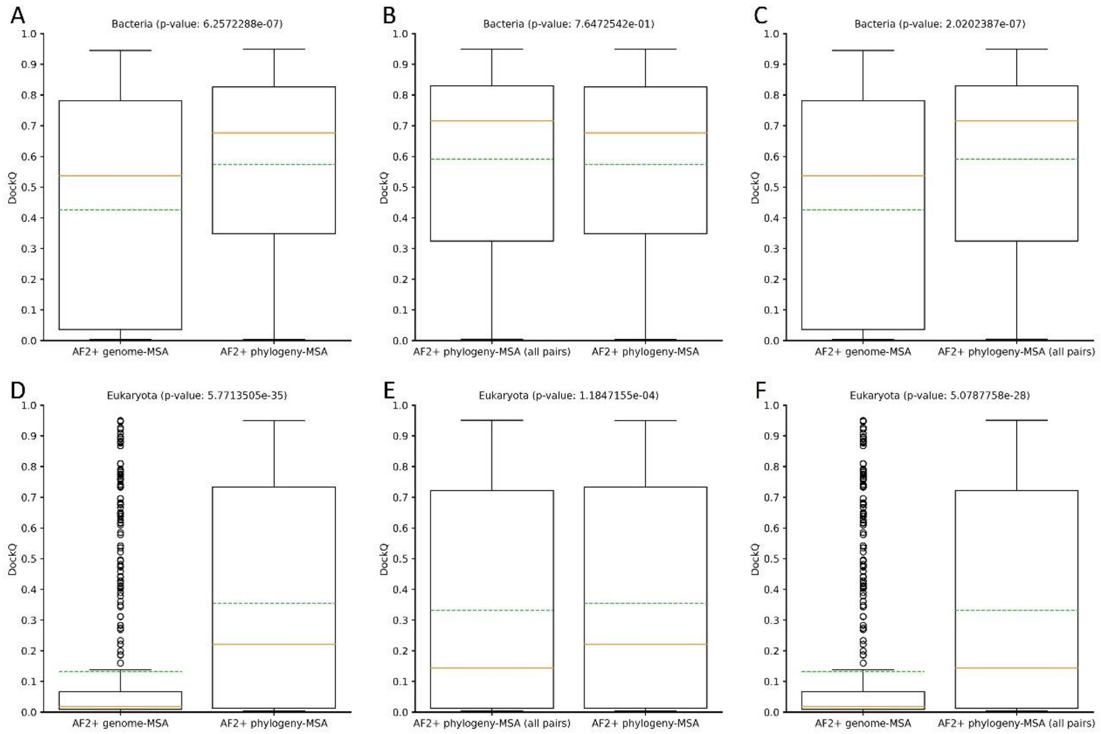


Figure S2. The performance comparisons between different protocols when using the Wilcoxon test to assess the differences. (A)~(C) the performance comparisons on the bacterial PPI dataset: (A) “AF2 + genome-MSA” versus “AF2 + phylogeny-MSA”; (B) “AF2 + phylogeny-MSA (all pairs)” versus “AF2 + phylogeny-MSA ; (C) “AF2 + genome-MSA” versus “AF2 + phylogeny-MSA (all pairs)”. (D)~(F) the performance comparisons on the eukaryotic PPI dataset: (D) “AF2 + genome-MSA” versus “AF2 + phylogeny-MSA”; (E) “AF2 + phylogeny-MSA (all pairs)” versus “AF2 + phylogeny-MSA ; (F) “AF2 + genome-MSA” versus “AF2 + phylogeny-MSA (all pairs)”. The protocols on the left were always used as the controls in the statistical tests.

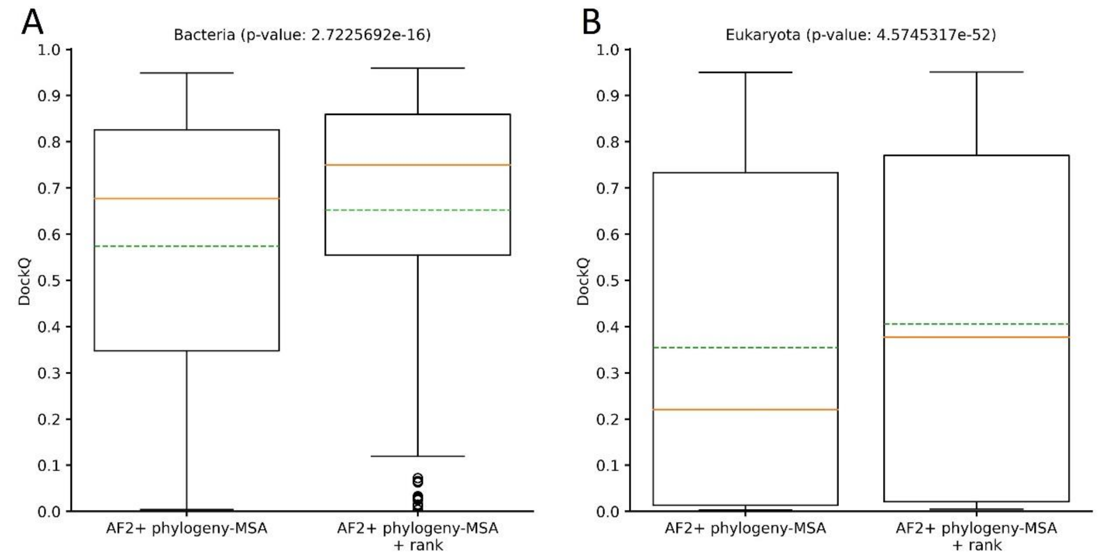


Figure S3. The performance comparisons between "AF2 + phylogeny-MSA" and "AF2 + phylogeny-MSA + rank" on (A) the bacterial PPI dataset and (B) the eukaryotic PPI dataset when using the Wilcoxon test to assess the differences. "AF2 + phylogeny-MSA" was used as the control in both statistical tests.

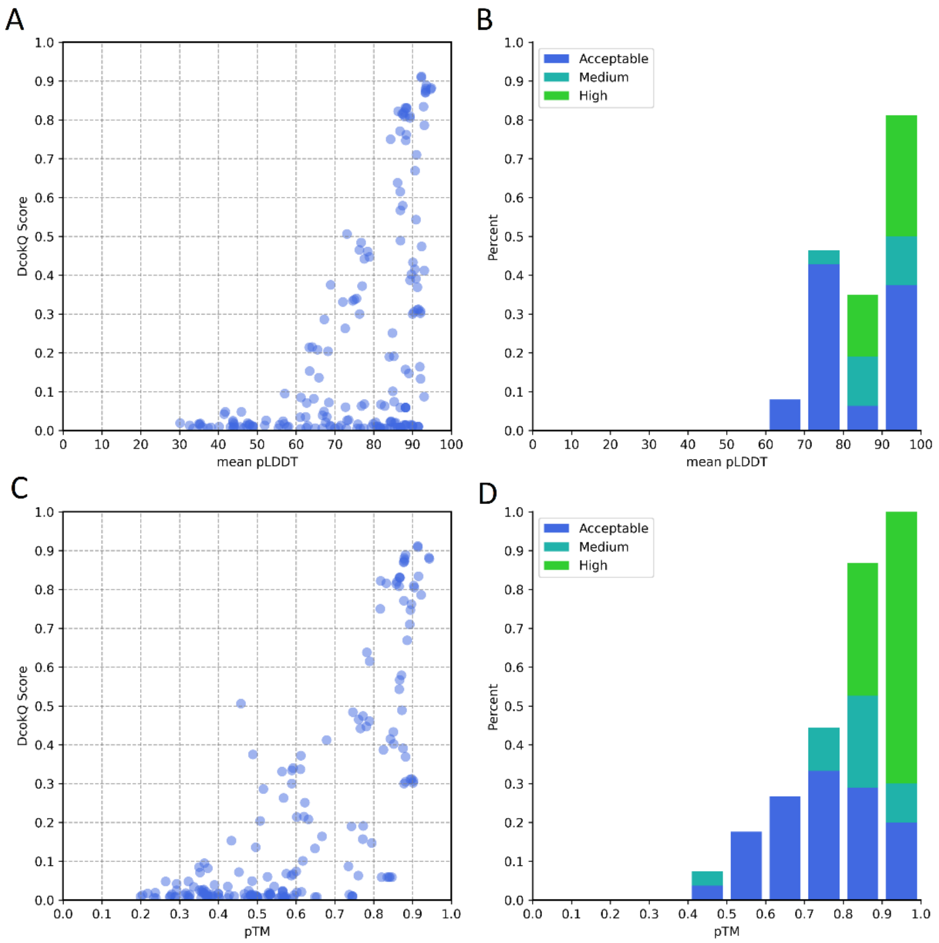


Figure S4. (A) DockQ versus mean pLDDT for each generated model, all the generated models using the MSAs of interologs from different taxonomic ranks of the species of the target PPIs for the 31 PPIs after AF2 training set are shown in the plot. (B) The proportions of the models with acceptable quality, medium quality and high quality when the mean pLDDT values are within different intervals. (C) DockQ versus pTM for each generated model, all the generated models using the MSAs of interologs from different taxonomic ranks of the species of the target PPIs for the 31 PPIs after AF2 training set are shown in the plot. (D) The proportions of the models with acceptable quality, medium quality and high quality when the pTM values are within different intervals.

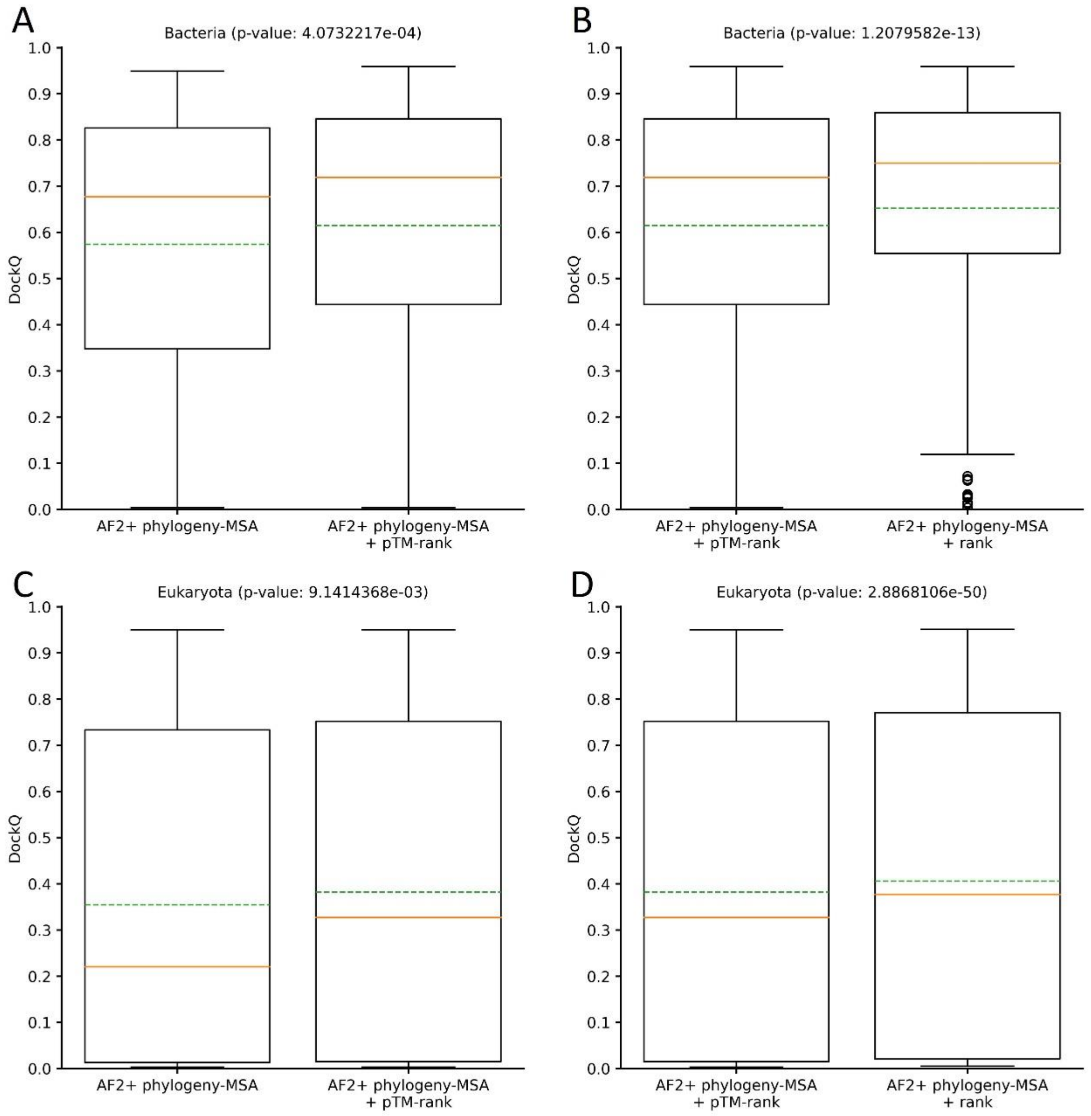


Figure S2. The performance comparisons between different protocols when using the Wilcoxon test to assess the differences. (A)~(B) the performance comparisons on the bacterial PPI dataset: (A) "AF2 + phylogeny-MSA" versus "AF2 + phylogeny-MSA + pTM-rank"; (B) "AF2 + phylogeny-MSA + pTM-rank" versus "AF2 + phylogeny-MSA + rank"; (C)~(D) the performance comparisons on the eukaryotic PPI dataset: (C) "AF2 + phylogeny-MSA" versus "AF2 + phylogeny-MSA + pTM-rank"; (D) "AF2 + phylogeny-MSA + pTM-rank" versus "AF2 + phylogeny-MSA + rank". The protocols on the left were always used as the controls in the statistical tests.

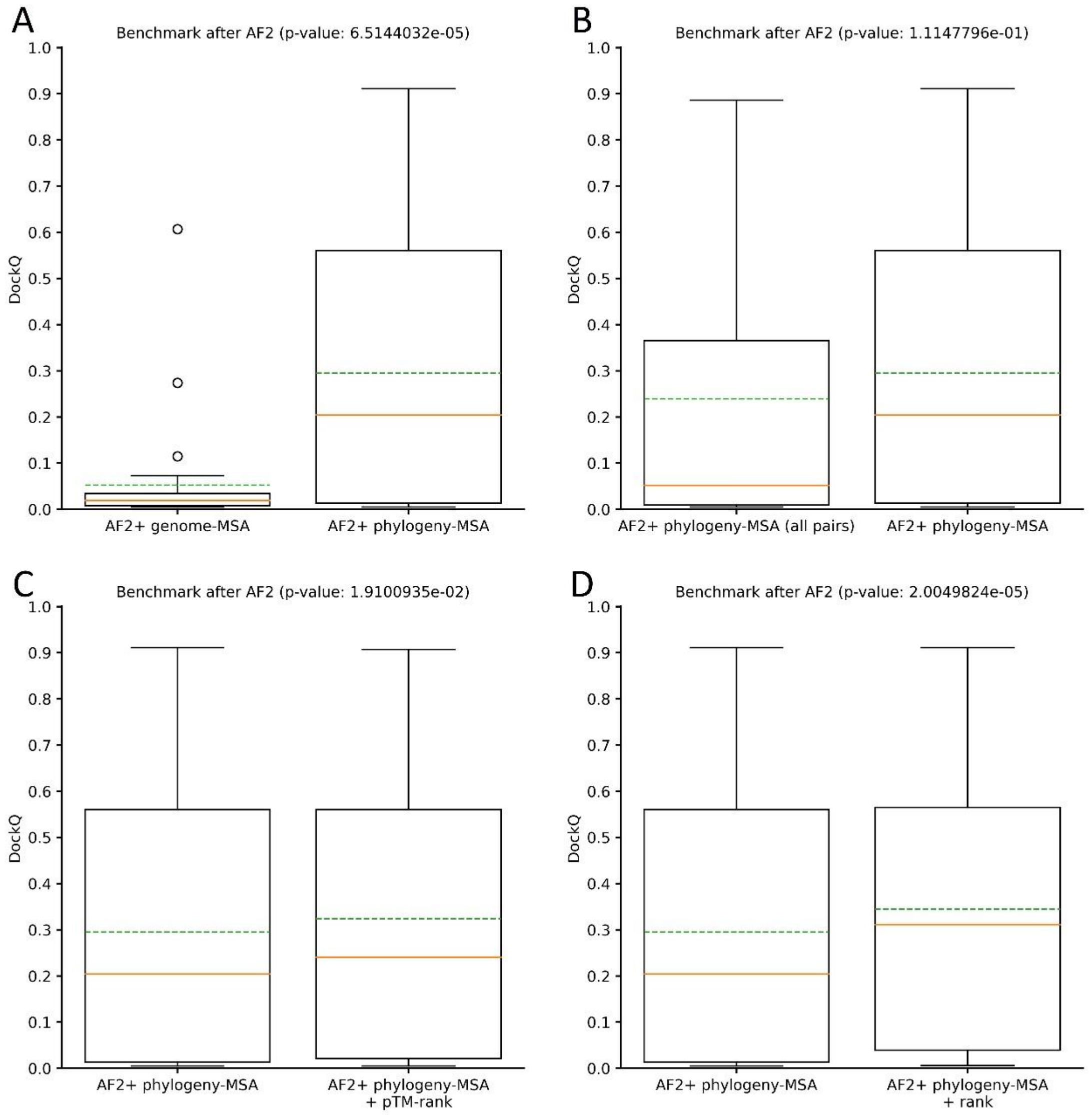


Figure S6. The performance comparisons between different protocols on the 31 PPIs after the training of AF2 when using the Wilcoxon test to assess the differences (A) “AF2 + genome-MSA” versus “AF2 + phylogeny-MSA”; (B) “AF2 + phylogeny-MSA (all pairs)” versus “AF2 + phylogeny-MSA” ; (C) “AF2 + phylogeny-MSA” versus “AF2 + phylogeny-MSA + pTM-rank”; (D) “AF2 + phylogeny-MSA” versus “AF2 + phylogeny-MSA + rank”; The protocols on the left were always used as the controls in the statistical tests.
